# Muscle-specific reticulospinal contributions to limb-trunk coordination during standing arm curls in strength-trained and untrained individuals

**DOI:** 10.64898/2026.07.18.739296

**Authors:** Nagisa Inubashiri, Shosuke Shinzaki, Hiroaki Kanehisa, Tadao Isaka, Sumiaki Maeo

## Abstract

Limb-trunk coordination plays an essential role in daily actions. The reticulospinal excitability of limb muscles has been suggested to be modulated by long-term motor experience, such as strength training. However, it remains unclear whether reticulospinal contributions in the limb and trunk muscles during limb-trunk coordinated movements are modulated by strength training. This study aimed to determine whether reticulospinal contributions to limb-trunk coordination differ between strength-trained and untrained individuals. Fifteen long-term (≥3 yrs) strength-trained and 15 untrained healthy men participated in this study. Participants performed a rapid bilateral arm-curl task while standing, using a load corresponding to 35% of their one-repetition maximum, in response to visual, visual-auditory (80 dB), or visual-startling (115 dB) stimuli. Electromyography (EMG) was recorded from the right biceps brachii (BB) and erector spinae (ES) muscles during the task. In the trained group, EMG onset of the ES was closer to that of the BB than in the untrained group, indicating tighter temporal coordination between the limb prime mover and trunk postural muscles in strength-trained individuals. In both groups, visual-startling stimuli shortened the EMG onset of both the BB and ES, suggesting reticulospinal contributions to both muscles. Notably, the reduction in EMG onset of the BB induced by the startling stimulus was smaller in the trained group than in the untrained group, whereas no difference between groups was observed for the ES. These findings suggest that long-term strength training may modify limb-trunk muscle coordination during standing arm curls and may be associated with muscle-specific adaptations in reticulospinal contributions.

## Introduction

Successful execution of limb movements during standing relies on the ability to maintain postural stability. A key mechanism underlying such limb-trunk coordination is anticipatory postural adjustments (APAs) of the trunk muscles, which involve the activation of trunk muscles that occurs in a period from −100 ms to +50 ms with respect to an intended limb movement [1–3]. APAs are altered by factors such as aging, neurological disorders, and motor learning [4–6]. The finding that motor learning can modify APAs indicates that APAs are not fixed processes. This finding also suggests that the neural control of APAs may differ between individuals with different motor-experience backgrounds. However, it remains unclear whether long-term motor experience induces neural adaptations not only in limb muscles but also in trunk muscles that maintain postural stability in response to perturbations caused by limb movements.

The corticospinal tract has been considered one of the major descending pathways involved in the neural control of APAs in trunk muscles. Previous studies have demonstrated that corticospinal excitability of the trunk muscles increases during the APAs and is modulated in a task- and age-dependent manner [7–10]. More recently, subcortical circuits, particularly the reticulospinal tract, have also been suggested to contribute to APAs in trunk muscles. Using the StartReact paradigm, an established indirect approach for assessing reticulospinal drive [11,12], several recent studies reported that the reticulospinal tract may contribute to APAs of back extensor muscles during rapid shoulder flexion [3,13]. To further understand the experience-dependent plasticity of neural circuits underlying APAs, it is essential to determine whether the reticulospinal contributions to limb-trunk coordination are modified by long-term motor experience.

Strength training may be an effective mode for modulating reticulospinal excitability. Recent evidence from a cross-sectional study suggests that strength training may enhance reticulospinal drive in upper-limb muscles [14]. Moreover, a non-human primate study has shown that short-term strength training increases reticulospinal excitability projecting to the trained muscles [15]. These findings suggest that reticulospinal pathways exhibit training-related plasticity. However, previous studies have primarily focused on muscles directly involved in the movement being trained. During many strength-training exercises performed in standing, force production by the upper limbs is accompanied by activation of trunk muscles to maintain postural stability [16]. Therefore, long-term strength training may induce not only neural adaptations in the prime mover muscles but also adaptations in the neural control of trunk muscles involved in limb-trunk coordination. Nevertheless, little is known about whether strength-trained individuals show altered reticulospinal contributions to trunk muscle activation during rapid upper-limb force production.

In the StartReact paradigm, differences in reticulospinal contributions between strength-trained and untrained individuals have been estimated based on the relative shortening of EMG onset induced by a startling auditory stimulus, referred to as RST gain. Previous studies examining movements involving the hand or elbow have reported inconsistent directions of strength training-related changes in RST gain [14,17]. These discrepancies may reflect differences in the relative contribution of the reticulospinal tract to force production, as reticulospinal drive is thought to be stronger in proximal and flexor muscles of the upper limb [18,19]. Accordingly, lower RST gain in upper-limb flexor muscles, such as the elbow flexors, in strength-trained individuals may reflect greater reticulospinal drive [14].

Therefore, the present study aimed to determine whether reticulospinal contributions to limb-trunk coordination during a rapid standing arm-curl task differ between strength-trained and untrained individuals. To this end, we compared temporal coordination between elbow flexor and trunk extensor activation, as well as the RST gain of these muscles, in strength-trained and untrained individuals. We hypothesized that strength-trained individuals would show altered temporal coordination between elbow flexor and trunk extensor activation and muscle-specific differences in RST gain compared with untrained individuals. Clarifying this issue would advance our understanding of experience-dependent modulation of subcortical contributions to limb-trunk coordination in healthy adults.

## Materials and Methods

### Participants

Thirty healthy men participated in this study, including 15 trained (age: 25.5 ± 5.8 years; height: 172.9 ± 6.4 cm; body mass: 75.7 ± 5.4 kg) and 15 untrained (age: 22.1 ± 1.5 years; height: 169.3 ± 5.1 cm; body mass: 58.8 ± 7.8 kg) individuals. Trained individuals were classified according to previously established criteria for long-term resistance-trained individuals [20]. Specifically, trained individuals had an extensive history of upper-arm resistance training with ≥ 2 sessions/week, ≥ 10 months/year, ≥ 3 years, isometric elbow flexion maximum voluntary torque of > 90 Nm, and maximum voluntary torque/body mass of > 1.1 Nm/kg. Untrained individuals had no experience with systematic resistance training. Participants provided written informed consent to participate in the study prior to the experiment. This study was conducted in accordance with the Declaration of Helsinki, and all procedures were approved by the Ethics Committee of Ritsumeikan University (Approval number: BKC-LSMH-2025-018).

### Experimental procedures

The experiment was conducted over two days. On the first day, the maximum voluntary torque was measured following a standardized warm-up protocol consisting of isometric voluntary contractions of 3-s duration at 50% (×3), 80% (×2) of perceived maximal effort. Participants were instructed to flex the elbow as hard as possible with verbal encouragement. On the second day, one-repetition maximum (1RM) for the bilateral arm-curl exercise was measured prior to the main experiment to determine the load used during the StartReact task. Participants then performed the StartReact task described below.

### StartReact task

Participants performed a rapid bilateral arm-curl task in response to audiovisual stimuli (Fig. 1). During the task, participants stood 1 m in front of a monitor with their feet shoulder-width apart while holding a barbell corresponding to 35% of their 1RM. Based on pilot testing, a load corresponding to 35% of 1RM was selected to allow participants to perform the arm-curl movement as rapidly as possible while providing sufficient postural demand to elicit APAs of the trunk muscles and minimizing fatigue during the experiment. Participants were instructed to flex both elbows to 90º as quickly as possible when a circular visual stimulus (diameter: 14 cm) appeared on the monitor. To assess the StartReact responses, three stimulus conditions were presented: visual-alone (V), visual-auditory (VA: 80 dB; 500 Hz; 50 ms), and visual-startling (VS: 115 dB; 500 Hz; 50 ms). The auditory stimulus was delivered from a speaker (IRX112BT, JBL, Northridge, USA) positioned at 0.3 m behind the participants. Decibel levels were confirmed using a digital sound level meter (30-130 dB: FN029A, YU-AN, Okinawa, Japan) before the task. The experimental task was implemented using a custom-written LabVIEW script (National Instruments, Texas, USA). Participants completed six blocks, each consisting of one practice trial followed by five trials of each stimulus condition presented in a pseudorandom order. Consequently, 30 trials were collected for each condition. The inter-trial interval was randomly varied between 4.5 and 6.5 s to prevent predictive movement initiation. Before data collection, participants performed 10 familiarization trials under the V condition. To minimize fatigue, a 3-min rest period was provided between the blocks.

**Figure 1.**
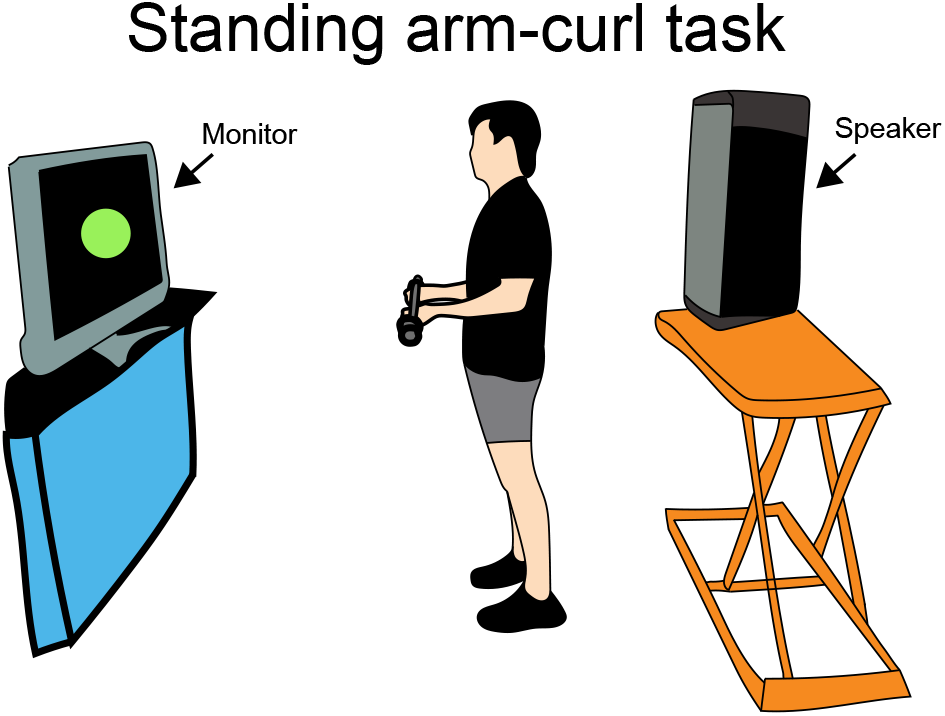
Experimental setup. Participants stood in front of a monitor with their feet shoulder-width apart while holding a barbell. Auditory stimuli were delivered by a speaker positioned behind the participants.

### Electromyography recording

Surface electromyograms (EMG) were recorded from the right biceps brachii long head (BB) and erector spinae (ES) muscles during the task. The EMG was recorded using bipolar Ag-AgCl electrodes (EM-272W, Noraxon, Arizona, USA). Each electrode had a diameter of 10 mm, and the interelectrode distance was 20 mm. The EMG electrode for the BB was placed according to SENIAM recommendations (seniam.org). The EMG electrode for the ES was placed 3 cm lateral to the T12 spinous process [3]. The skin was prepared by shaving the hair, surface abrading, and cleansing with 80% ethanol. Reference electrodes were placed over the bony prominence of the wrist on both limbs. Each electrode was connected to an amplifier (MEG-6108MMG, Miyuki Giken, Japan), where the EMG signal was band-pass filtered between 5 and 1000 Hz and amplified (×1000). All electrical signals were recorded at a sampling frequency of 2000 Hz and stored on the hard disk of a personal computer using a 16-bit analogue-to-digital converter (cDAQ-9187, National Instruments, Texas, USA).

### Data analysis

The EMG signal was analyzed using a custom-written MATLAB script (R2025a, The MathWorks, Massachusetts, USA). The raw EMG signals were band-pass filtered between 20 and 500 Hz using a zero phase-lag fourth-order Butterworth filter, demeaned, and rectified [13]. The onset time of the EMG was identified as the first point at which the rectified EMG exceeded three standard deviations above the mean background EMG level, calculated from the 50-ms period preceding the stimulus, and remained for at least 20 ms. The EMG onset was searched within 1000 ms following stimulus onset. All onset detections were visually inspected while blinded to the stimulus condition to confirm the automatically detected onset times. Preliminary analyses indicated that EMG onset times gradually decreased during the initial trials and reached a plateau during the latter half of the experiment (see Supplementary Figure 1). Therefore, only the final 15 trials from each condition were included in subsequent analyses to minimize the influence of task familiarization. The EMG onset times were averaged across the 15 trials within each condition. To quantify the temporal coordination between BB and ES activation, the relative ES onsets were calculated by subtracting the EMG onset of BB from that of ES. Finally, to compare the reticulospinal excitability between groups, the RST gain was calculated as ([V–VS]/[V–VA]) [12,14,17], where larger values indicate a greater shortening of onset time in response to the startling stimulus relative to the non-startling auditory stimulus.

### Statistical analysis

We confirmed the normality of all the tested data by using Shapiro-Wilk tests (*p* > 0.05). We performed two-way repeated-measures analysis of variance (ANOVA) to investigate differences in relative ES onsets among stimulus conditions and between the trained and untrained groups. Also, the differences in EMG onsets among stimulus conditions and between the trained and untrained groups were tested using two-way repeated-measures ANOVA. Significant ANOVA results were followed up by post hoc multiple comparisons using the Bonferroni correction. We performed Student’s *t*-test to compare muscle strength (absolute maximum voluntary torque and relative maximum voluntary torque) and RST gain between the trained and untrained groups. All statistical analyses were conducted using MATLAB (R2025a, The MathWorks, Massachusetts, USA) and JASP (version 0.97.1, JASP Team, Amsterdam, The Netherlands) [21] with a significance level of 0.05.

## Results

### Muscle strength

To confirm that the trained and untrained groups represented distinct training backgrounds, absolute maximum voluntary torque, body-weight-normalized maximum voluntary torque, and 1RM were compared between groups. The trained group had 80% higher absolute maximum voluntary torque (98.8 ± 6.5 vs. 54.8 ± 8.4 Nm, *p* < 0.001), 39% higher relative maximum voluntary torque (1.31 ± 0.11 vs. 0.94 ± 0.13 Nm/kg, *p* < 0.001), and 119% higher 1RM (48.47 ± 6.04 vs. 22.10 ± 4.10 kg, *p* < 0.001) than the untrained group.

### Temporal coordination of BB and ES onsets

Figures 2A and 2B show the distributions of BB and ES onsets in the trained and untrained groups, respectively. We observed that the differences between BB and ES onsets in trials were consistent among stimulus conditions (Fig. 2C). Two-way repeated-measures ANOVA revealed no main effects of stimulus condition (*F*2,56 = 2.00, *p* = 0.145) and the group × stimulus interaction (*F*_2,56_ = 1.06, *p* = 0.354). In contrast, a significant main effect of group was observed (*F*_1,28_ = 5.64, *p* = 0.025), indicating that ES onset occurred closer in time to BB onset in the trained group than in the untrained group.

**Figure 2.**
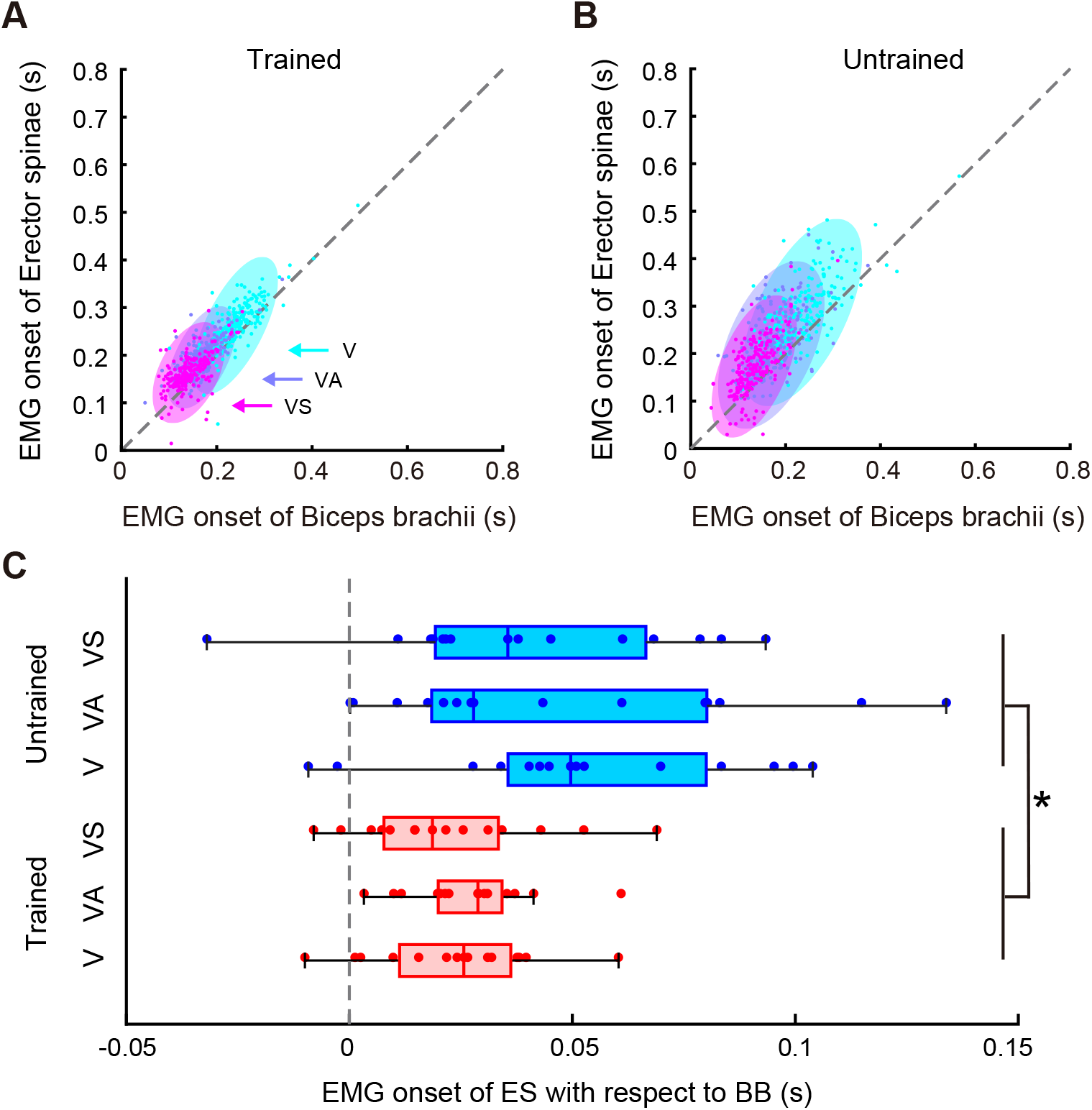
Temporal relationship of EMG onsets between the biceps brachii and the erector spinae. **A**: Distribution of the EMG onsets of the biceps brachii and the erector spinae in the trained group. **B**: Distribution of the EMG onsets of the biceps brachii and the erector spinae in the untrained group. The dots represent the EMG onsets in each trial, pooled across all analyzed data of all participants. Ellipses indicate 95% confidence intervals. The cyan, deep sky blue, and magenta dots and ellipses correspond to the visual (V), visual-auditory (VA), and visual-startling (VS) conditions, respectively. **C**: EMG onset of the erector spinae with respect to the biceps brachii in the trained (red) and untrained (blue) group. Lines indicate median values and outer edges of the box represent the 25th and 75th percentiles. The upper and lower whiskers indicate maximum and minimum values. Each dot represents the mean value in each condition for an individual participant. The asterisk indicates *p* < 0.05.

### StartReact effect

Fig. 3A illustrates the rectified EMG traces of the BB and ES during the V, VA, and VS conditions in a representative trained participant. The mean EMG onsets for the BB and ES are presented in Fig. 3B. For the BB, a two-way repeated-measures ANOVA revealed a significant main effect of stimulus condition (*F*_2,56_ = 327.2, *p* < 0.001). In contrast, neither the main effect of group (*F*_1,28_ = 0.25, *p* = 0.624) nor the group × stimulus interaction (*F*^2,56^ = 0.54, *p* = 0.585) was significant. Post hoc analyses revealed that BB onset in both groups was significantly shorter in the VA and VS conditions than in the V condition and was further shortened in the VS condition compared with the VA condition (*p* < 0.001). Similar results were observed for the ES. Two-way repeated-measures ANOVA revealed a significant main effect of stimulus condition (*F*_2,56_ = 225.4, *p* < 0.001), whereas neither the main effect of group (*F*_1,28_ = 2.52, *p* = 0.124) nor the group × stimulus interaction (*F*_2,56_ = 1.26, *p* = 0.290) was significant. Post hoc analyses showed that ES onset in both groups was significantly shorter in the VA and VS conditions than in the V condition and was further shortened in the VS condition relative to the VA condition (*p* < 0.001).

**Figure 3.**
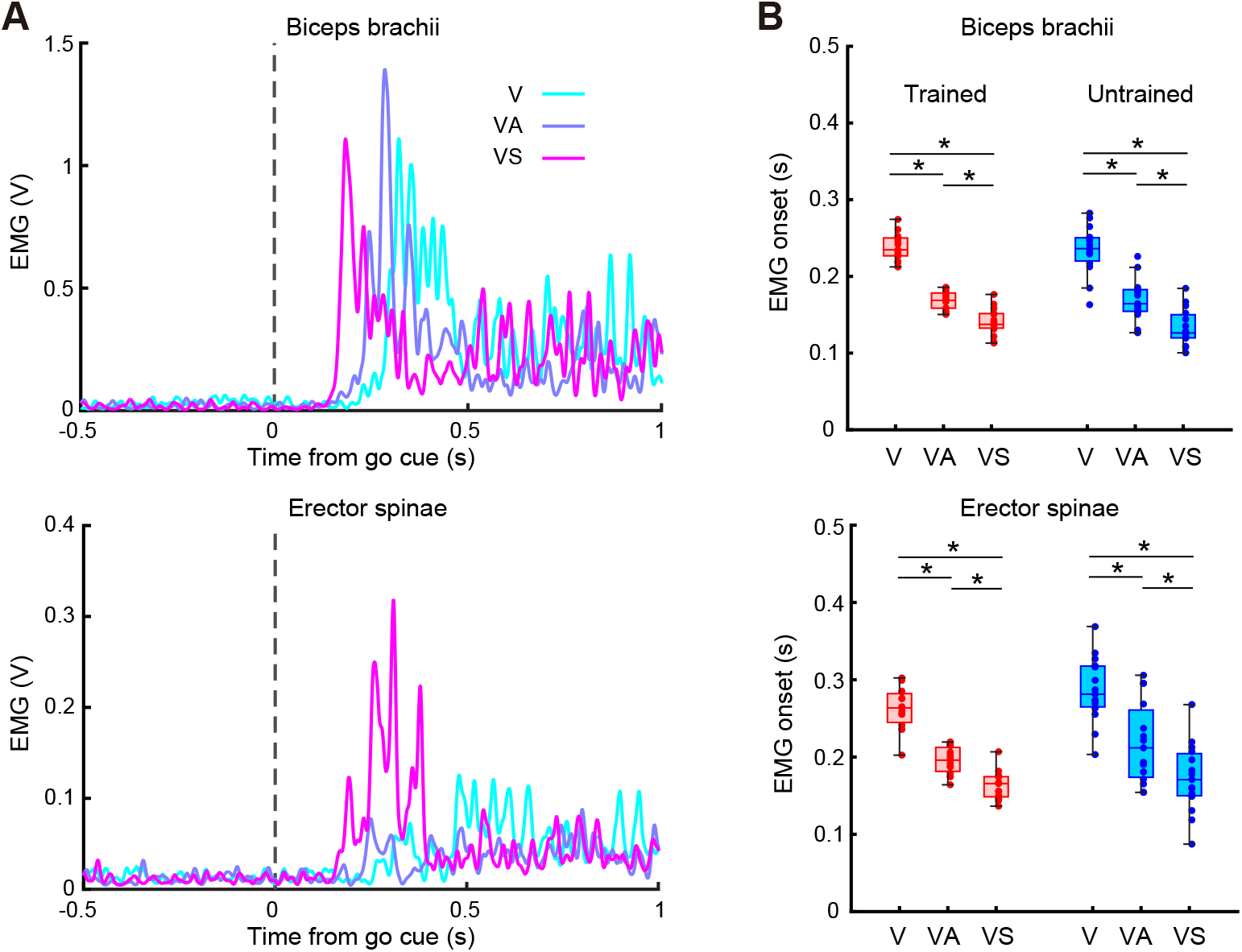
StartReact responses in the biceps brachii and the erector spinae. **A:** EMG traces of the biceps brachii (upper panel) and the erector spinae (lower panel) in the visual (V), visual-auditory (VA), and visual-startling (VS) conditions of a representative participant. The dotted line indicates the stimulus onset. **B**: EMG onsets of the biceps brachii (upper panel) and the erector spinae (lower panel) in the trained (red) and untrained (blue) group. Lines indicate median values and outer edges of the box represent the 25th and 75th percentiles. The upper and lower whiskers indicate maximum and minimum values. Each dot represents the mean value in each condition for an individual participant. The asterisk indicates *p* < 0.05.

### RST gain

To examine group differences in reticulospinal excitability, the RST gain was compared between groups. Fig. 4 shows the mean RST gain of the trained and untrained groups. For the BB, the trained group showed a significantly lower RST gain than the untrained group (*p* = 0.019: Fig. 4A). In contrast, no significant group difference was observed in the RST gain of the ES (*p* = 0.165: Fig. 4B).

**Figure 4.**
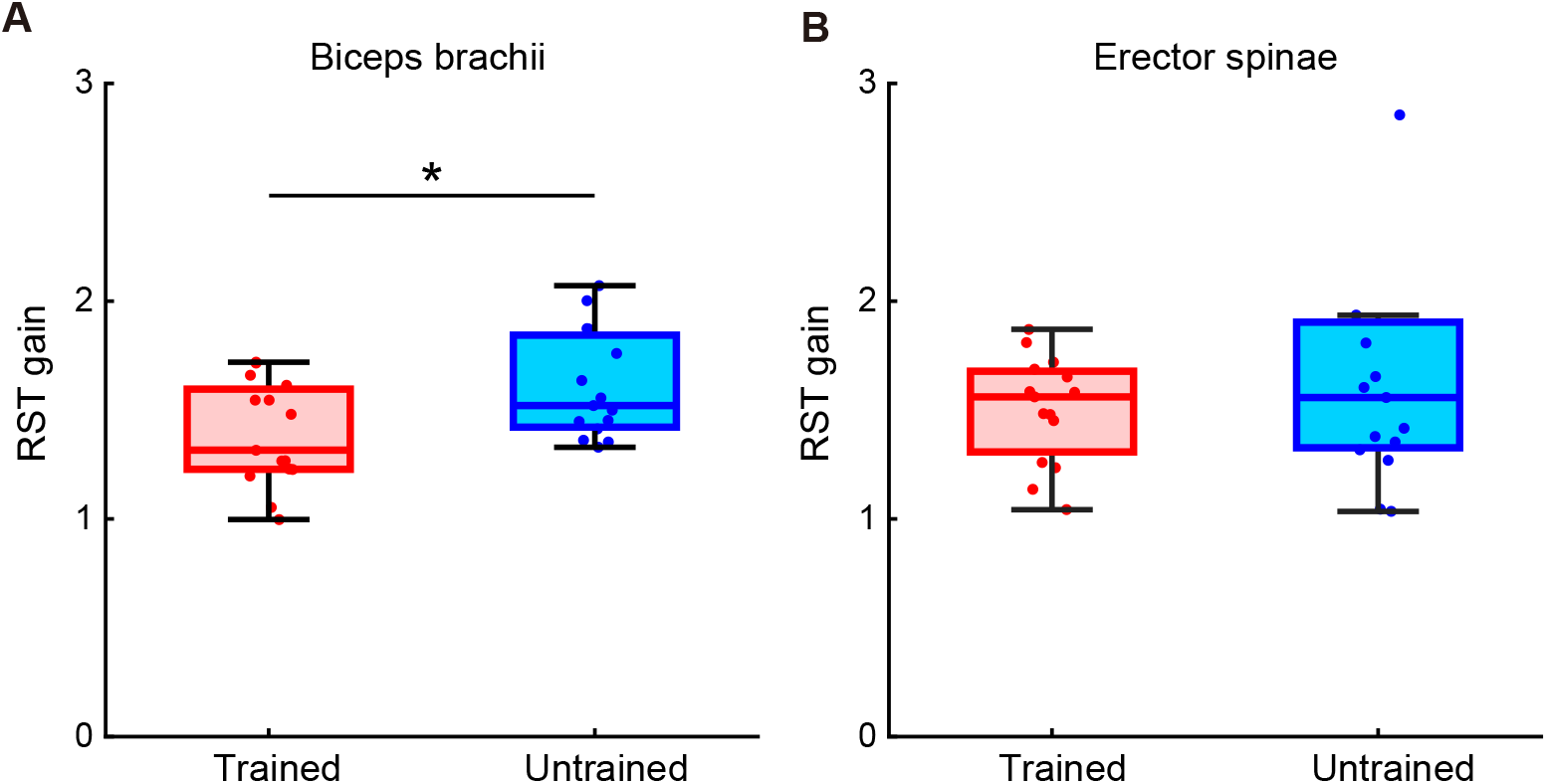
RST gain. **A**: RST gain of the biceps brachii in the trained (red) and untrained (blue) group. **B**: RST gain of the erector spinae in the trained (red) and untrained (blue) group. Lines indicate median values and outer edges of the box represent the 25th and 75th percentiles. The upper and lower whiskers indicate maximum and minimum values. Each dot represents the mean value for an individual participant. The asterisk indicates *p* < 0.05.

## Discussion

In this study, we aimed to investigate whether reticulospinal contributions in the limb and trunk muscles during a rapid bilateral elbow flexion while standing differ between strength-trained and untrained individuals. To address this issue, we used the StartReact paradigm in which participants performed the bilateral arm-curl movements in response to visual, non-startling auditory, and startling auditory stimuli. In the trained group, the timing of APAs of the ES was closer to the EMG onset of the BB than in the untrained group. StartReact effects were present in both the BB and ES muscles in both groups, suggesting that the startling auditory stimulus accelerated the release of a prepared motor command for arm-curl movement, probably through subcortical pathways involving the reticulospinal tract. However, the trained group showed a lower RST gain in the BB, but not in the ES, compared with the untrained group. These findings suggest that strength-trained individuals show altered reticulospinal excitability in the prime mover (i.e., BB), but not in the trunk muscle, together with altered temporal coordination between the limb and trunk.

Our results show that APA-related ES activation occurred closer in time to BB onset in the trained group than in the untrained group. These findings suggest that strength training enhances temporal coordination between the limb prime mover and trunk postural muscles in healthy adults. In line with this study, previous studies have reported that APA timing in patients with musculoskeletal or neurological disorders can be advanced by short-term strength training [22–24]. In addition, in healthy young adults, dynamic perturbation training has been shown to induce earlier anticipatory muscle activations during a predictable postural control task [25]. Therefore, the earlier APA timing observed in strength-trained individuals in the present study may share common features with training-related changes in APA timing reported in previous studies. To our knowledge, this study is the first to demonstrate tighter temporal coordination between prime mover and trunk muscle activation in healthy strength-trained individuals than in untrained individuals.

Despite the tighter temporal coordination between BB and ES activation in trained individuals, RST gain in the ES did not differ significantly between the trained and untrained groups. In both groups, EMG onsets of the ES were shortened by the presentation of a startling auditory stimulus, suggesting that the reticulospinal tract contributes to ES activation during the standing arm-curl task [26]. However, APAs are believed to be mediated by distributed cortical, subcortical, and brainstem networks, including the temporoparietal and parietal cortices, supplementary and premotor areas, cerebellum, basal ganglia, and corticoreticulospinal pathways [27]. Taken together, these findings suggest that the training-induced adaptation of ES activation timing during upper-limb movements may involve neural mechanisms other than an increase in reticulospinal drive, at least as assessed by the RST gain.

The RST gain in BB was lower in the trained group than in the untrained group, whereas the RST gain in ES was comparable between groups. Consistent with previous studies [3,13], StartReact effects were observed in both the BB and ES during the limb-trunk coordinated movements, suggesting that the reticulospinal tract contributes to the activation of both the prime mover and the trunk postural muscle. To extend our understanding of plasticity of the reticulospinal tract involved in limb-trunk coordination, we investigated whether long-term strength training experience influences reticulospinal excitability of the elbow flexor and the trunk extensor during arm-curl movements. The lower RST gain observed in the trained group compared with the untrained group, but only in the BB, suggests that strength training may induce muscle-specific adaptations in reticulospinal contributions. During the standing arm-curl movement, the BB acts as a prime mover, whereas the ES acts as a postural muscle. Given that previous research has shown that reticulospinal pathways may contribute to ES activity regardless of whether the muscle acts as a prime mover or as a postural muscle [13], long-term strength training may preferentially modify reticulospinal contributions to muscles that are repeatedly recruited as prime movers during the trained movement, rather than uniformly enhancing reticulospinal drive to all muscles involved in limb-trunk coordination.

As discussed above, the direction of training-related differences in RST gain may depend on the relative contribution of the reticulospinal tract to the target muscle [14,17–19]. Because elbow flexors are thought to receive relatively strong reticulospinal input, the lower RST gain of the BB in the trained group may reflect enhanced reticulospinal drive even in the absence of a startling auditory stimulus [14,18,19]. In contrast, the absence of a group difference in ES RST gain suggests that long-term strength training may not uniformly alter reticulospinal contributions to all muscles involved in limb-trunk coordination. Thus, the present findings support the idea that strength training-related differences in reticulospinal contribution are muscle-specific, at least in the context of standing arm-curl movements.

We investigated the differences in reticulospinal excitability between the trained and untrained individuals. It should be noted that the present study cannot determine the neural substrates underlying the altered APA timing in the trained group. In addition, because the current study was cross-sectional, we cannot clarify a causal relationship between long-term strength training and the observed differences in APA timing or reticulospinal excitability. Therefore, training intervention studies focusing on a broad range of neural systems would help clarify whether strength training directly induces these adaptations and identify the underlying neural mechanisms.

## Conclusion

In this study, we demonstrated that ES activation occurred closer in time to BB onset during arm-curl movements in the trained group than in the untrained group. Furthermore, RST gain of the elbow flexor was lower in the trained group than in the untrained group, whereas such a difference was not observed in the trunk extensor. These results suggest that long-term strength training may modify APAs during upper-limb force production and may induce muscle-specific adaptation of the reticulospinal tract. These findings enhance our understanding of the plasticity of the neural circuits responsible for limb-trunk coordinated movements, emphasizing the effect of strength training on reticulospinal excitability.

## CRediT authorship contribution statement

**Nagisa Inubashiri**: Conceptualization, Funding acquisition, Methodology, Investigation, Formal analysis, Data curation, Project administration, Software, Visualization, Writing – original draft. **Shosuke Shinzaki**: Investigation. **Hiroaki Kanehisa**: Writing – review & editing. **Tadao Isaka**: Funding acquisition, Supervision. **Sumiaki Maeo**: Conceptualization, Funding acquisition, Supervision, Writing – review & editing.

## Funding

This work was supported by a Research grant from Tobe Maki Foundation to N.I; JSPS KAKENHI Grant Number 25K24332 to N.I.; JSPS KAKENHI Grant Number JP25K03016 to S.M.; and JSPS Program for Forming Japan’s Peak Research Universities (J-PEAKS) Grant Number JPJS00420240020 to N.I. and T.I.

## Declaration of competing interest

The authors report no conflict of interest.

## Supplementary Figure

**Supplementary Figure 1.**
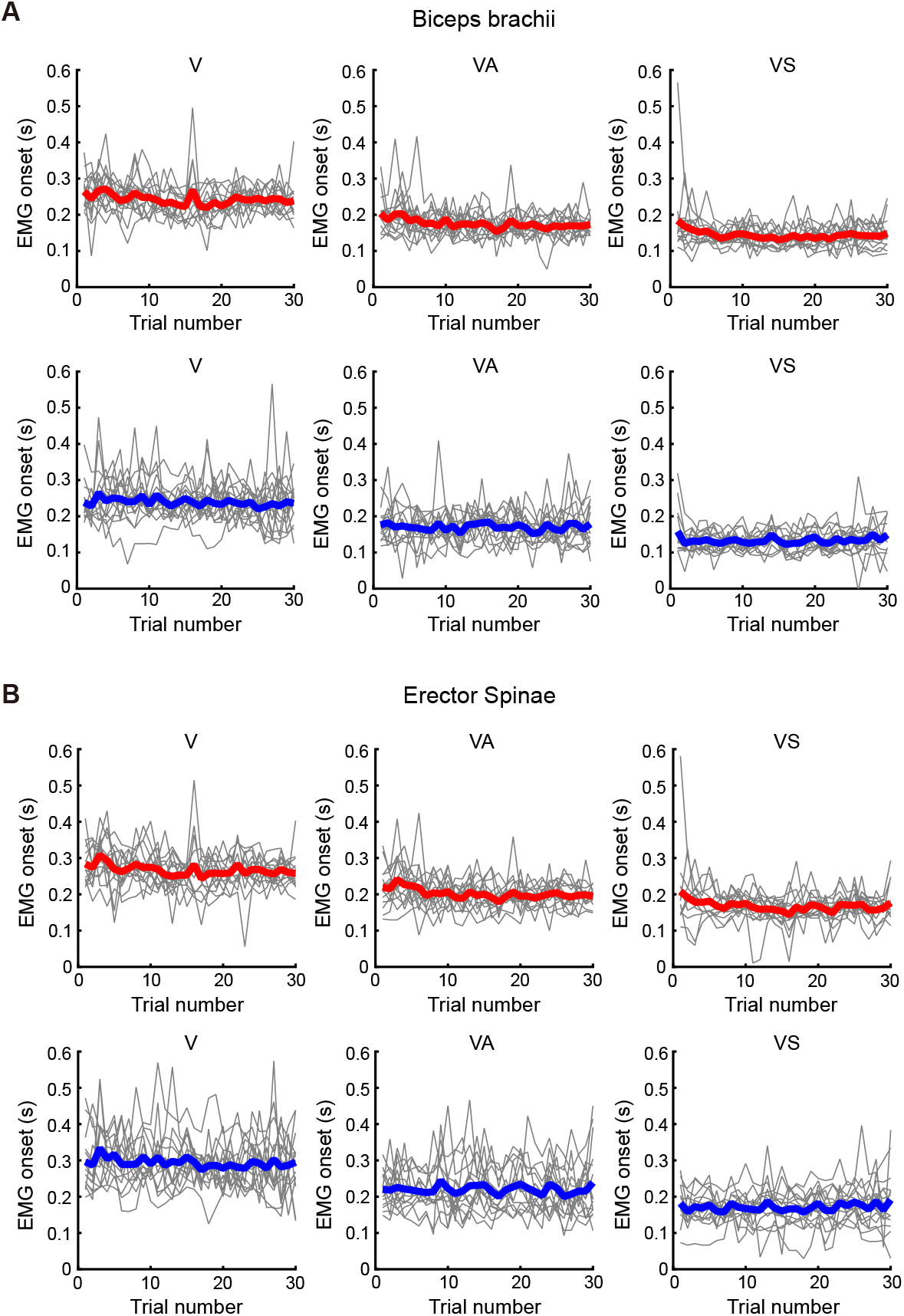
Time course of the EMG onset in the visual (V), visual-auditory (VA), and visual-startling (VS) conditions during the experiment. **A**: EMG onsets of the biceps brachii in the trained (upper panel) and untrained (lower panel) group. **B**: EMG onsets of the erector spinae in the trained (upper panel) and untrained (lower panel) group. The red and blue lines indicate the average across participants in the trained and untrained groups, respectively. The gray lines represent individual participants.

## Notes

### Competing Interest Statement

The authors have declared no competing interest.

